# Androgen signaling connects short isoform production to breakpoint formation at Ewing sarcoma breakpoint region 1 via an R-loop-dependent mechanism

**DOI:** 10.1101/2020.03.25.008391

**Authors:** Taylor R. Nicholas, Peter C. Hollenhorst

## Abstract

Ewing sarcoma breakpoint region 1 (*EWSR1*) encodes a multifunctional protein that can cooperate with the transcription factor ERG to promote prostate cancer. The EWSR1 gene is also commonly involved in oncogenic gene rearrangements in Ewing sarcoma. Despite the cancer relevance of *EWSR1*, its regulation is poorly understood. Here we find that in prostate cancer, androgen signaling upregulates a 5’ *EWSR1* isoform by promoting usage of an intronic polyadenylation site. This isoform encodes a cytoplasmic protein that can strongly promote cell migration and clonogenic growth. Deletion of an Androgen Receptor (AR) binding site near the 5’ *EWSR1* polyadenylation site abolished androgen-dependent upregulation. This polyadenylation site is also near the Ewing sarcoma breakpoint hotspot, and androgen signaling promoted R-loop and breakpoint formation. RNase H overexpression reduced breakage and 5’ *EWSR1* isoform expression suggesting an R-loop dependent mechanism. These data suggest that androgen signaling can promote R-loops internal to the *EWSR1* gene leading to early transcription termination and breakpoint formation.

## Introduction

Ewing’s sarcoma breakpoint region 1 (*EWSR1*) is ubiquitously expressed in humans and plays an essential role in normal development (Azuma et al., 2007; Li et al., 2007; Yamamoto-Shiraishi et al., 2014). In both Ewing’s sarcoma and certain leukemias, oncogenic gene rearrangements can fuse the 5’ end of *EWSR1* to the 3’ end of transcription factor encoding genes. One such fusion, *EWSR1/FLI1*, typifies Ewing’s sarcoma as it is found in 85% of cases (Grunewald et al., 2018). The fusion protein, EWS/FLI1, binds the genome through the C-terminal DNA binding domain of FLI1, while the N-terminal EWS portion functions as a strong transactivation domain (TAD) (Rossow and Janknecht, 2001). Because of the outstanding recurrence of EWS/FLI1 in Ewing’s sarcoma, most of our understanding of EWS function comes from the fusion context. However, the wild-type EWS protein clearly has multivariate roles that are not well understood.

Both nuclear and cytoplasmic (Andersson et al., 2008), EWS, the protein encoded by *EWSR1*, has transcription-dependent and transcription-independent roles. We have previously reported that EWS can act as a transcriptional co-activator in prostate cancer (Kedage et al., 2016); other reported nuclear functions of EWS include the regulation of splicing and DNA damage repair (Dutertre et al., 2010; Gorthi et al., 2018; Paronetto et al., 2011). Sedimentation studies suggest that cytoplasmic EWS associates with dense, ribosome-containing fractions as well as lighter fractions that also contain the plasma membrane (Felsch et al., 1999). Additionally, knockout studies have implicated *EWSR1* in meiosis, with null mice defective in spermatogenesis and oogenesis (Li et al., 2007).

EWS belongs to a small family of proteins with conserved structure and partially overlapping functions, termed the FET family. FET family members have an N-terminal prion-like domain (PrLD) consisting of degenerate SYGQ repeats. This “low complexity domain” has important transcription-related oncogenic properties as the N-terminus of all three FET proteins are found fused in various cancers to transcription factor genes (Grunewald et al., 2018; Nyquist et al., 2011; Riggi et al., 2007). In the C-terminus, FET proteins bind nucleic acids with an RNA recognition motif (RRM) and a zinc finger domain (ZnF), punctuated by arginine-glycine-glycine rich (RGG) regions. At the very C-terminal end is a nuclear localization signal. Aside from cancer, all three FET family members are mutated in ALS and in frontotemporal dementia (FTD) (Couthouis et al., 2012; Kapeli et al., 2017). FET proteins have been demonstrated to form higher order structures that cause liquid demixing and formation of membraneless organelles through aggregation in the PrLD (Altmeyer et al., 2015; Qamar et al., 2018; Tanikawa et al., 2018) or RNA binding (Maharana et al., 2018). When reversible, these “droplets” facilitate normal cellular processes such as cytoplasmic stress granule formation but when irreversible may form pathological plaques in disease (Patel et al., 2015).

Androgen receptor (AR) is a ligand-inducible nuclear hormone receptor that is important in development and disease. Once activated by androgens, AR translocates to the nucleus and functions as a transcription factor, controlling genetic programs important for growth and tissue determination (Huang et al., 2014). AR promotes male-specific characteristics and is involved in prostate cancer initiation and progression (Lonergan and Tindall, 2011; Murashima et al., 2015). In addition to transcriptional function, AR also promotes the chromosomal rearrangement responsible for the *TMPRSS2/ERG* gene fusion found in 50% of prostate tumors (Haffner et al., 2010; Lin et al., 2009). Interestingly, like *EWSR1* null mice, *AR* null mice are defective in spermatogenesis from meiotic arrest (De Gendt et al., 2004), suggesting that *AR* and *EWSR1* can regulate similar phenotypes.

In this study, we find that androgen signaling regulates the *EWSR1* gene to produce different genetic outcomes important for cancer biology. In prostate cancer, we found that AR drives formation of a shortened *EWSR1* isoform that promotes cancer-associated phenotypes. We then used prostate cancer cells as a model to show androgen driven formation of a break in *EWSR1* at the same breakpoint hotspot that creates the *EWSR1/FLI1* oncogene in Ewing sarcoma. While it is known that AR promotes gene fusion formation in prostate cancer, an androgen-dependent mechanism for *EWSR1* breakage has not been shown. This is the first study to characterize direct androgen regulation of the *EWSR1* gene.

## Results

### Androgen signaling upregulates an intronic polyadenylated *EWSR1* isoform

We previously found that EWS plays an important role in prostate cancer (Kedage et al., 2016), therefore we investigated the relationship between *EWSR1* expression and *AR* expression, as AR is essential for prostate cancer development and progression. Using the prostate adenocarcinoma (PRAD) TCGA dataset housed on UCSC Xena Browser (https://xenabrowser.net) (Goldman, 2018), *EWSR1* mRNA level was ranked by *AR* gene expression in 550 patient tumor samples. Mean expression of all *EWSR1* exons negatively correlated with *AR* expression (Figure S1A). However, exon-level analysis found that the 5’ *EWSR1* exons were positively correlated with *AR* while the 3’ exons were inversely correlated (Figure 1A). Xena Browser by default normalizes the log2(RPKM+1) values for each exon. To determine absolute expression levels, patient samples were separated into two bins: top 25% (high AR) and bottom 25% (low AR) and mean expression of each exon was plotted (Figure 1B, left). The 5’ *EWSR1* exons showed the greatest separation of signal between the two bins, with the exceptions of exon 1 (a 5’ UTR exon), exon 5a (a brain specific exon), and exons 9-11 (which Xena browser called as a single exon including the intervening introns). Exons 12 and 13 remained at a similar expression level, regardless of AR expression, while the 3’ most exons were expressed at lower levels when AR was high. Plotting the difference of the two bins (Figure 1B, right) reveals a pattern of progressively decreased exon expression across the EWSR1 gene when AR levels are high. Together, these data indicate that AR expression correlates with higher levels of 5’ EWSR1 exons and suggests that AR might promote early termination of the EWSR1 transcript. To understand the specificity of this observation, we performed the same analysis across multiple cancer types using the PANCAN TCGA data set (Figure 1C). In this case, only a modest (log2(RPKM) = ∼0.2) difference in 5’ exon expression was seen, however the decrease in 3’ exons still occurred.

**Figure 1.**
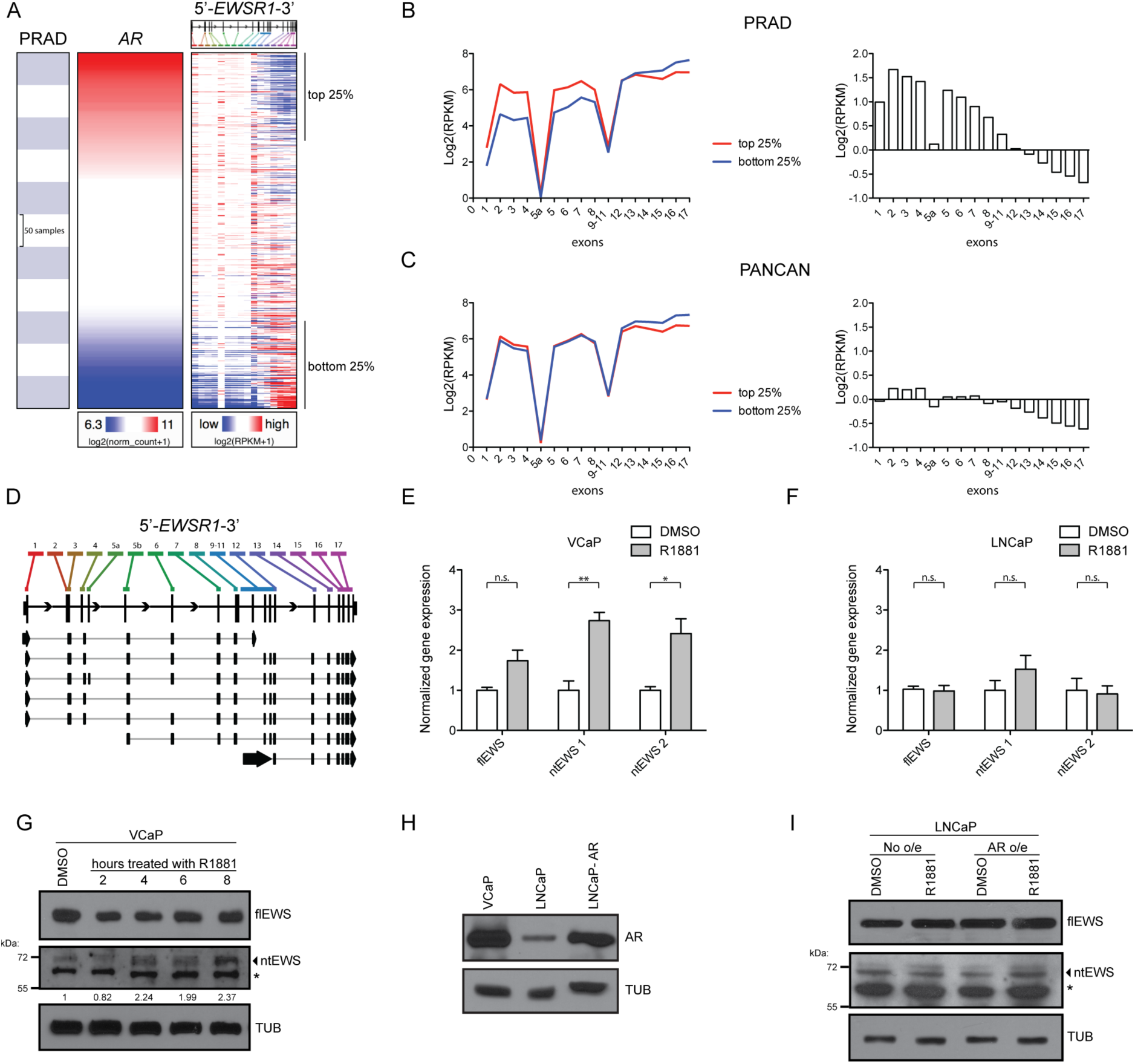
Androgen signaling upregulates an intronic polyadenylated *EWSR1* isoform. A) *EWSR1* exon expression ranked by *AR* gene expression in primary prostate cancer patients (PRAD). B) (right) Average expression of *EWSR1* exons in the samples with the highest 25% of *AR* expression (red) and with the lowest 25% of *AR* expression (blue) for the PRAD data set. (left) Difference in expression of the top 25% expression bin and the bottom 25% expression bin. C) Same as in B) but for the PANCAN TCGA data set. D) UCSC Xena Browser called exons for *EWSR1* (top) with gene schematic (middle) and hg19 annotated *EWSR1* isoforms (bottom). E-F) Normalized gene expression by isoform-specific qRTPCR for *flEWS* and *ntEWS* measured by one or two isoform-specific primer sets, respectively, in VCaP (E) and LNCaP (F) cells. Expression is normalized to 18S and relative to the DMSO condition. Shown is mean ± SEM for three replicates. All p values (*p < 0.05, ** p < 0.01) were obtained by t tests. G) Immunoblot of ntEWS in VCaP treated with 10 nM R1881 for indicated time. Quantification of ntEWS bands are normalized to tubulin (TUB) and relative to DMSO. H) AR immunoblot in indicated cell lines. I) Immunoblot of flEWS, ntEWS, and TUB in LNCaP cells with or without AR overexpression treated with 10 nM R1881 for 6 hours. See also Figure S1.

The differing expression level of the 5’ and 3’ *EWSR1* exons suggests that *EWSR1* might not be transcribed as a single unit and that multiple *EWSR1* isoforms exist. The hg19 UCSC gene annotation (Haeussler et al., 2019) for *EWSR1* shows several isoforms (Figure 1D), including one (NM_001163287.2) that is composed of 5’ exons and terminates via an intronic polyadenylation event that generates an alternative last exon (ALE), exon 9, and 3’ UTR. Because this isoform encodes the N-terminus of the EWS protein, we have termed it the N-terminal isoform or ntEWS. Although annotated, *ntEWS* regulation and function has not been reported in the literature to our knowledge. Polya_db (Wang et al., 2018) (version 3.2, http://exon.njms.rutgers.edu/polya_db/v3/), a database of polyA sites (PAS), captures the intronic polyadenylation event that forms *ntEWS* in 69.2% of samples (Figures S1B and S1C). Genomic 3’-sequencing data (Lianoglou et al., 2013) indicates that ntEWS is preferentially expressed in the testis (Figure S1D), consistent with a male-specific and possibly androgen-regulated role.

Since all exons in *ntEWS* track with AR levels in patient tumors (besides exon 1 and 9, mentioned above), we tested whether androgen signaling would increase *ntEWS* RNA levels. Androgen responsive prostate cancer cell lines VCaP and LNCaP were treated with 10 nM synthetic androgen R1881 for 24 hr and *ntEWS* mRNA measured by qRTPCR. Primers specific for *ntEWS* span the novel exon-exon junction to the ALE and *flEWS* specific primers test the first exon not present in *ntEWS*. Prostate specific antigen (PSA) RNA was measured to verify androgen response (Figure S1E). Treatment of VCaP cells with R1881 caused a significant upregulation in *ntEWS* mRNA level measured by two distinct primer sets. Full-length *EWS* expression increased modestly but not to a significant level (Figure 1E). In contrast, treatment with R1881 did not increase the level of *ntEWS* in LNCaP cells (Figure 1F).

To measure levels of the ntEWS protein, a polyclonal rabbit antibody was generated using the sequence encoded by the ALE as an epitope. The ∼70 kDa band was verified as ntEWS by the presence of a similar migrating band in PC3 cell lysates overexpressing *ntEWS* (Figure S1F), and by diminished levels of this band upon treatment with three independent *ntEWS* shRNAs (Figure S1G). To determine if ntEWS protein increased after androgen exposure, VCaP cells were treated for 2, 4, 6, or 8 hours with R1881 and ntEWS protein levels were compared to the DMSO control (Figure 1G). After 4 hours of R1881, ntEWS protein increased approximately two-fold (Figure 1G).

We questioned whether the ability of R1881 to increase *ntEWS* levels in VCaP, but not LNCaP was due to differences in AR expression levels. In fact, AR was more abundant in VCaP than in LNCaP (Figure 1H), which corresponded to heightened AR activity in VCaP measured by PSA expression (Figure S1E). To test the role of AR expression, AR was overexpressed in LNCaP cells (Figure 1H) and ntEWS protein and PSA RNA were measured after treatment with R1881. AR overexpression in LNCaP cells allowed R1881 to promote a further increase in PSA (Figure S1E), and, strikingly, mediated an R1881-dependent upregulation of ntEWS protein (Figure 1I).

### AR binding to Intron 5 of *EWSR1* directly regulates ntEWS expression

To determine if AR regulates the *EWSR1* gene through direct binding, we analyzed published AR ChIP-seq datasets from patient tumor samples and matched adjacent normal tissue (Pomerantz et al., 2015). AR binds to two intragenic *EWSR1* sites; one in intron 5, and one near the exon-intron boundary of exon 9 and intron 8 that overlaps with the PAS that produces ntEWS (Figure 2A). AR occupancy at the intron 5 site was particularly strong in four of the six tumors and was higher than any matched normal peak, suggesting tumor-specific binding of AR to this site. Interestingly, the patients with a higher tumor:normal signal at the intron 5 site have later staged, more aggressive disease compared to the patients with a low tumor:normal signal, measured by pathological staging and biochemical recurrence (Pomerantz et al., 2015). These four patient’s tumors appear to have generally higher AR binding as they have high AR enrichment at the well characterized AREs near *KLK2* and *KLK3* (Figure S2A).

**Figure 2.**
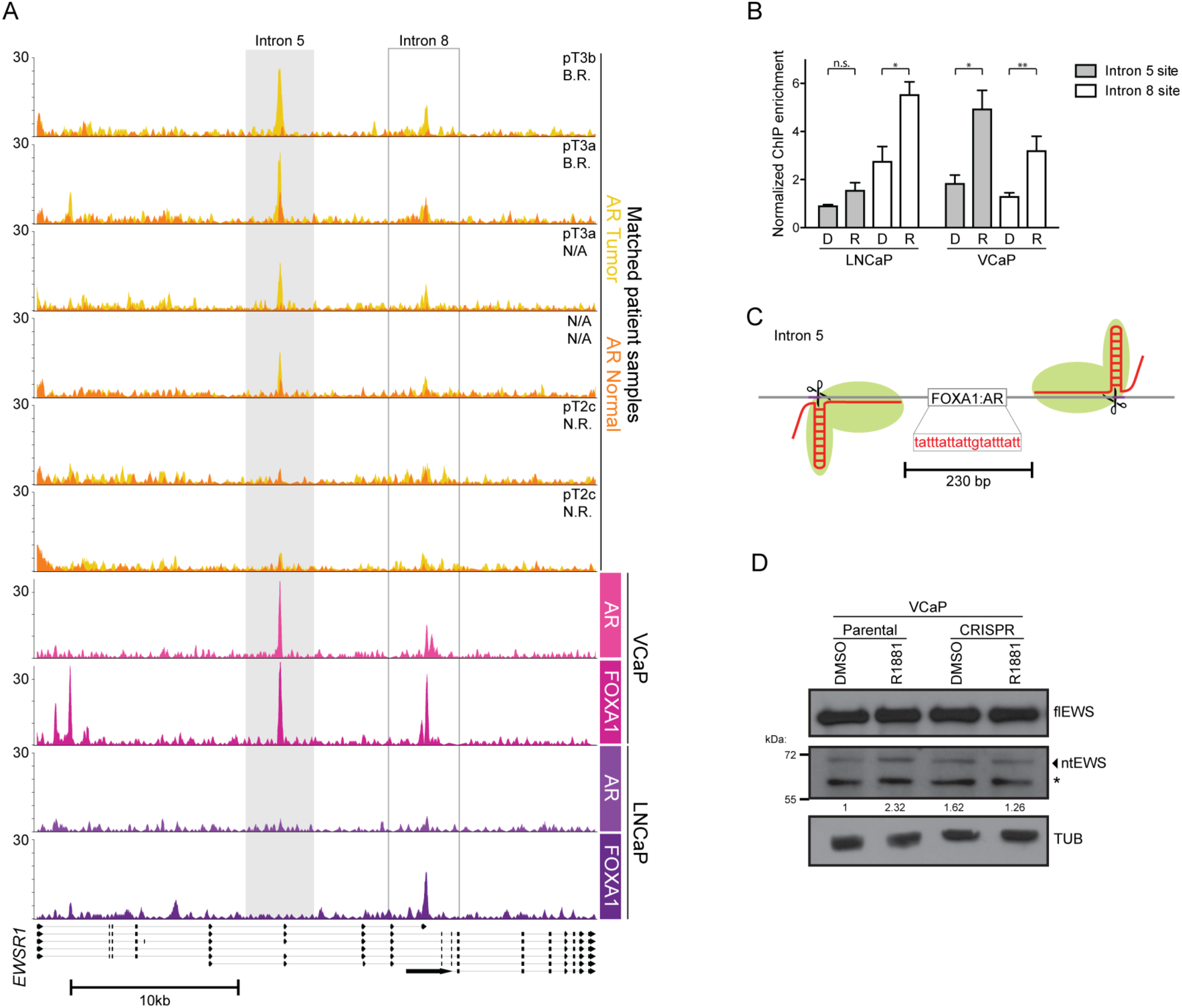
AR binding to Intron 5 of *EWSR1* directly regulates ntEWS expression. A) Gene tracks for ChIP-seq of AR or FOXA1 in samples as labeled. Pathological stage and biochemical recurrence (B.R.) or no recurrence (N.R.) is indicated for tumors on the top right. B) ChIP-qPCR of AR in LNCaP and VCaP cells treated with DMSO (D) or R1881 (R) at the Intron 5 (gray) and Intron 8 (white) *EWSR1* sites. ChIP enrichment is normalized to a negative control region and the mean ± SEM for three replicates is shown. p values (*p < 0.05, ** p < 0.01) were obtained by t tests. C) Depiction of the pgRNA targeting strategy for the FOXA1:AR binding site in intron 5 of *EWSR1*. D) Immunoblot of flEWS, ntEWS or tubulin (TUB) in parental or CRISPR VCaP cells treated with DMSO or 10 nM R1881 for 6 hr. Quantification as in Figure 1G. See also Figure S2.

AR ChIP-seq in VCaP cells treated with 10 nM R1881 for 2hrs (Toropainen et al., 2015) shows similar localization observed in patient tumors at the EWSR1 gene: modest signal at the intron 8 and robust signal at the upstream intron 5 site (Figure 2A). However, in LNCaP cells treated with 100 nM DHT for 2hrs (Malinen et al., 2017), AR shows very low levels of binding to both sites (Figure 2A), suggesting that high levels of AR protein is required to saturate these binding sites. The AR co-factor FOXA1 was bound to both sites in VCaP cells and to the intron 8 site only in LNCaP cells. To reconcile treatment differences and verify binding, we performed ChIP-qPCR of AR in VCaP and LNCaP cells treated with 10 nM R1881 for 2hrs. AR was significantly enriched at both sites when stimulated with R1881 in VCaP cells in a manner consistent with the ChIP-seq data (Figure 2B). In LNCaP cells, AR only showed significant binding to the intron 8 site (Figure 2B).

The sequence at each AR binding site identified in the ChIP-seq data was analyzed to identify androgen response elements (AREs). Three different AR position weight matrices (JASPAR) were tested using the FIMO tool from MEME-suite (Grant et al., 2011). The most significant AR motif was a FOXA1:AR site in intron 5 (p = 5.06e-05). Since AR binding at the intron 5 site is more robust in tumors and VCaP cells, we focused on understanding the importance of AR binding to that site by a CRISPR-Cas9 disruption strategy. To avoid the limitations of targeting a short A:T rich sequence, paired guide RNAs (pgRNA) were targeted to sequence flanking the intron 5 ARE (Figure 2C) as pgRNA targeting is an effective method to achieve targeted deletions (Diao et al., 2017; Gasperini et al., 2017; Thomas et al., 2020). Cells with Cas9 targeted to the intron 5 ARE failed to upregulate ntEWS upon R1881 treatment while not affecting flEWS expression (Figure 2D). This suggests that the ARE in intron 5 is an important and specific *cis*-regulatory element for *ntEWS*.

### ntEWS promotes phenotypes related to oncogenesis

Since the function of ntEWS has not been described in the literature, we cloned and overexpressed ntEWS in the androgen insensitive cell line PC3. For comparison, we also overexpressed flEWS and a predicted C-terminal isoform (ctEWS) that shares no sequence homology to ntEWS (Figure S3A). Nuclear-cytoplasmic fractionation of the over expressing lines showed the presence of both flEWS and ctEWS in the nucleus and cytoplasm; however, ntEWS was found exclusively in the cytoplasm (Figure 3A), consistent with previous reports of the nuclear localization signal in the EWS C-terminus (Shaw et al., 2009).

**Figure 3.**
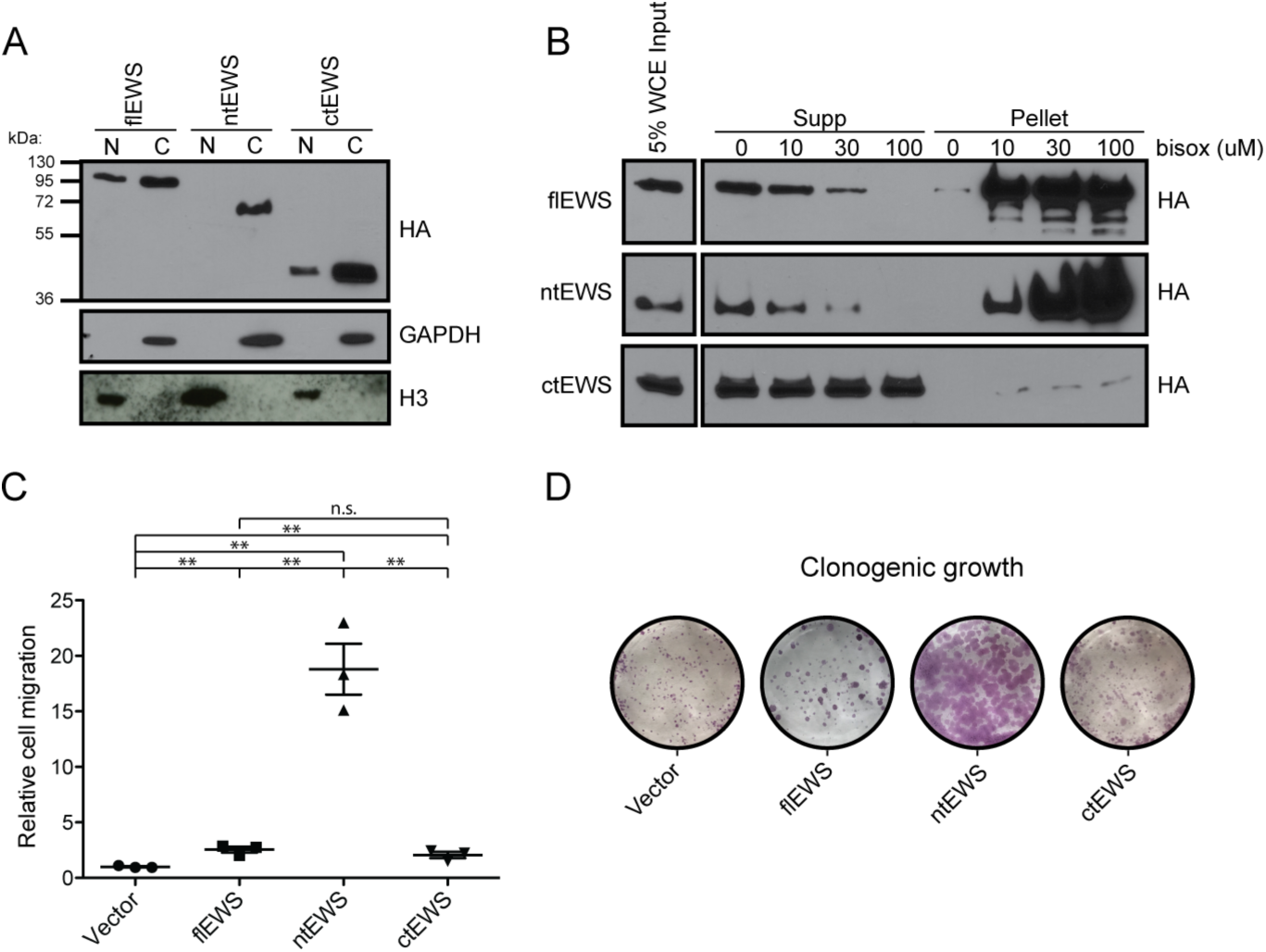
ntEWS promotes phenotypes related to oncogenesis. A) Nuclear-cytoplasmic fractionation of 3xHA-flEWS, 3xHA-ntEWS, and 3xHA-ctEWS in PC3 cells visualized by HA immunoblot. Nuclear and cytoplasmic fraction controls are H3 and GAPDH immunoblot, respectively. B) B-isox precipitation assay of PC3 with 3xHA-flEWS, 3xHA-ntEWS, or 3xHA-ctEWS immunoblotted for HA. C) Cell migration of EWS isoform overexpressing PC3 cells. Cell migration relative to the vector expressing cells is the mean ± SEM for three biological replicates with two technical replicates each. All p values (**p > 0.01) were obtained by t test. D) Clonogenic growth of EWS isoform overexpressing PC3 cells. Representative images are shown; two additional biological replicates were similar. See also Figure S3.

The N-terminal PrLD of EWS has been shown to form higher order structures seeded by various substrates in the context of flEWS and EWS/FLI1 (Boulay et al., 2017; Kato et al., 2012). To determine the ability of ntEWS to form such structures, we performed a biotinylated isoxazole (b-isox) precipitation assay. Addition of the compound b-isox to cell lysates seeds precipitates formed by low-complexity domains like those found in the PrLD of EWS (Kato et al., 2012). We added increasing amounts of b-isox to PC3 cell lysates containing flEWS, ntEWS or ctEWS. Consistent with previous reports (Kato et al., 2012), flEWS was found in the b-isox seeded pellet in a concentration dependent manner. ntEWS was also pelleted by b-isox in a concentration dependent manner, while ctEWS, which lacks the PrLD, was not precipitated by b-isox, even at high concentrations (Figure 3C).

We next sought to determine the cellular impact of EWS isoform over-expression on PC3 cells. Transwell cell migration and clonogenic growth assays were performed in overexpression lines. Expression of flEWS induced significant cell migration and clonogenic growth compared to vector expressing cells, consistent with our previous work (Kedage et al., 2016). Additionally, ctEWS expression promoted these phenotypes to a similar extent as flEWS. Strikingly, ntEWS expression promoted dramatic increases in cell migration and clonogenic growth (Figure 3D and 3E). ntEWS expressing cells did not show increased proliferation in normal growth conditions compared to the flEWS and ctEWS expressing cells, measured by MTT assay (Figure S3B), suggesting that ntEWS driven phenotypes are not simply a function of increased proliferation.

### The ntEWS alternative last exon encodes an alpha helical domain important for function

To gain insight on the structure of ntEWS compared to the N-terminus of flEWS, we compared ntEWS to EWS (1-355aa), which is the same length but lacks the ALE (Figure 4A). The online tool IUPRED (Dosztanyi, 2018) predicted disorder to be high across the EWS N-terminus, but low in the ALE (Figure S4A). Phyre2 (Kelley et al., 2015) was used to predict secondary structure. While the entirety of EWS (1-355aa) is predicted to be intrinsically disordered with high confidence, the 30aa ALE encoding region of ntEWS is predicted to be alpha helical with high confidence (Figure S4B). These data suggest that the unique region encoded by the ALE in the ntEWS isoform is structured unlike the rest of the N-terminus of EWS.

**Figure 4.**
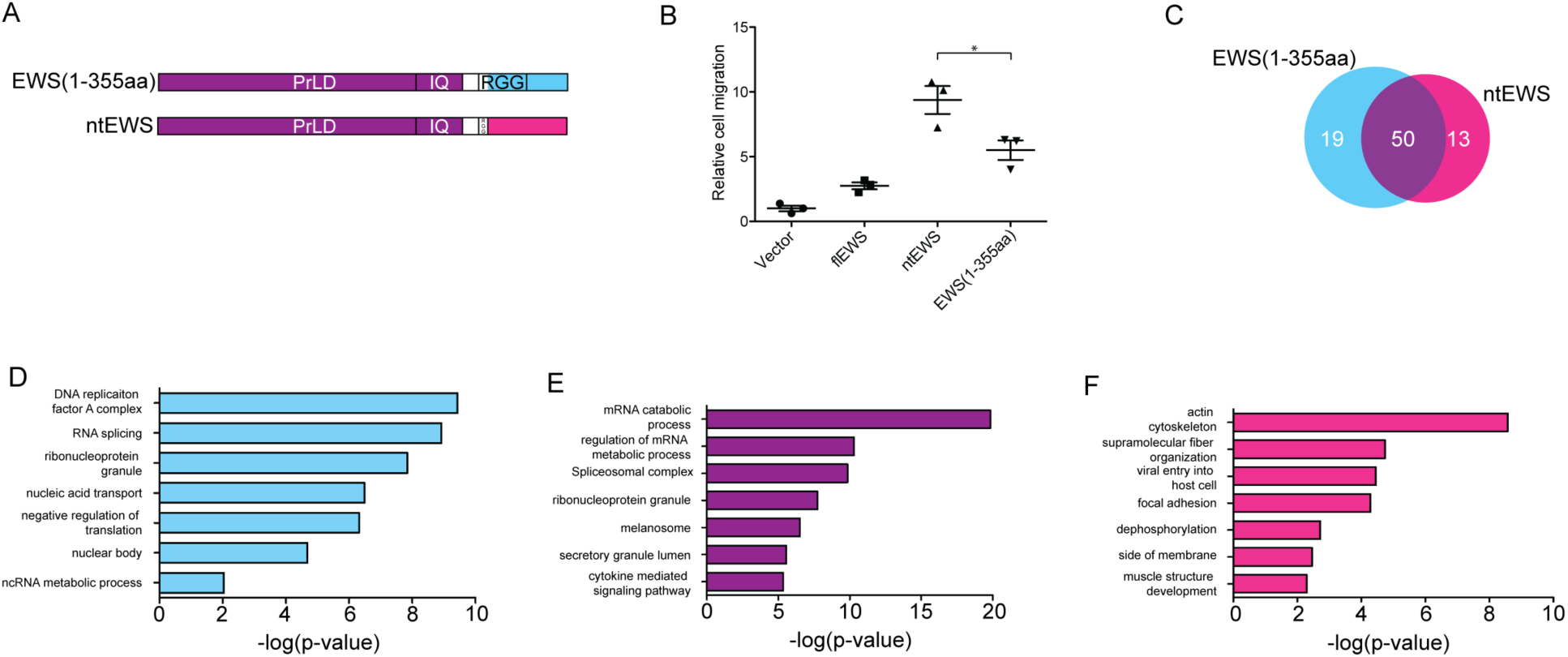
The ntEWS alternative last exon encodes an alpha helical domain important for function. A) Schematic diagram of EWS (1-355aa) and ntEWS. The Prion-like domain (PrLD), Calmodulin binding domain (IQ) and arginine glycine rich regions (RGG) regions are shown. B) Cell migration of 3xHA-ntEWS and 3xHA-EWS(1-355aa) overexpressing PC3 cells. Cell migration relative to vector is the mean ± SEM for three biological replicates with two technical replicates each. p value (*p < 0.05) were obtained by t tests. C) Venn diagram showing the number of interacting proteins in each category identified in two biological replicates of IP-MS. D-F) Biological process and cellular component gene ontologies of interacting partners. The top eight ontologies are shown as a function of the –log(p value). See also Figure S4.

To determine the functional contribution of the PrLD and the alpha helical domain of ntEWS, we performed functional assays in PC3 cells expressing HA-tagged ntEWS or EWS (1-355aa). Interestingly, EWS (1-355aa) induced cell migration, 6-fold more than the vector expressing cells, however not to the same extent as ntEWS, which drove cell migration 10-fold more than the control (Figure 4B). This suggests that the PrLD can promote cell migration alone, but the alpha helical domain of ntEWS contributes to the robust phenotype only observed in the ntEWS expressing cells.

To identify unique protein interactions with the ntEWS alpha helical domain, both ntEWS and EWS (1-355aa) were immunoprecipitated from the cytoplasm of PC3 cells and subjected to mass spectrometry in two independent replicates. To achieve a high confidence interactomes, only proteins that were called in both replicates and scored a 99.9% probability threshold were considered for analysis. We identified 19 proteins specific to EWS (1-355aa), 13 proteins for ntEWS, and 50 proteins as shared interacting partners (Figure 4C, Table 2) attributed to the common N-terminal PrLD. Many previously identified interaction partners of flEWS (Pahlich et al., 2009) were identified in “shared” category (16/50 proteins), including FET family members FUS and TAF15. Additionally, Calmodulin, which interacts with the IQ domain in the PrLD (Deloulme et al., 1997) was pulled down by both proteins. Gene ontology analysis of each category was performed using Metascape (Zhou et al., 2019) and the top eight significant ontologies related to biological processes and cellular components were plotted (Figure 5D, 5E, and 5F). The top ontologies for ntEWS specific interactors include actin cytoskeleton, supramolecular fiber organization, and focal adhesion (Figure 5F), consistent with a cytoplasmic role for ntEWS in cell migration, possibility through regulation of the actin cytoskeleton.

**Figure 5.**
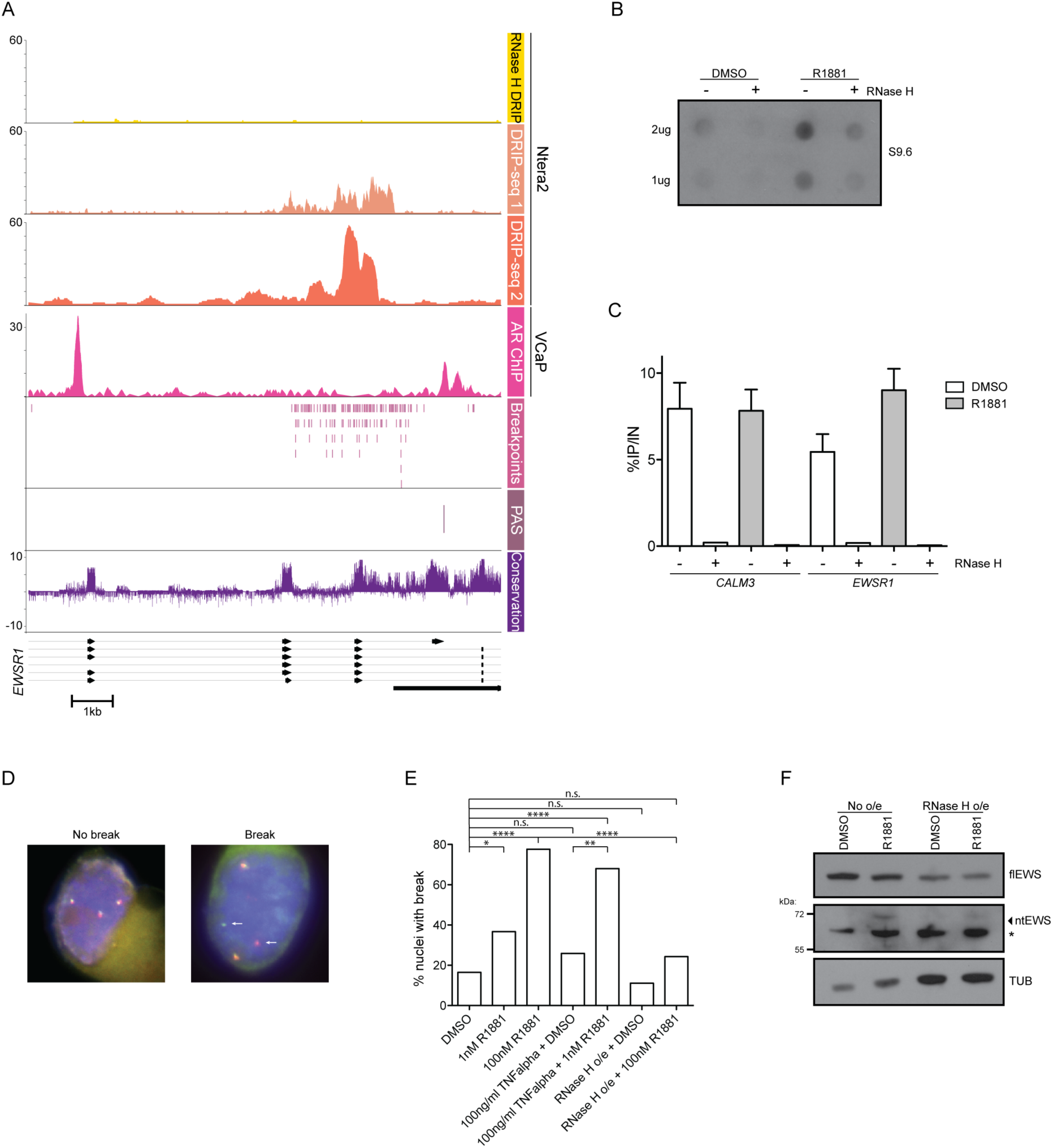
Androgen signaling promotes *EWSR1* breakpoint formation via R-loops. A) DRIP-seq enrichment from (Sanz et al., 2016) shown from exon 6 to exon 10 of *EWSR1*. Breakpoint coordinates in *EWSR1* are from the Catalog of Somatic Mutations In Cancer (COSMIC) (Tate et al., 2019). Conservation is from multiple alignment of 100 vertebrate species. B) Dot blot using DNA/RNA hybrid antibody for the labeled conditions. C) DRIP-qPCR for *CALM3* and intron 5 of *EWSR1* from VCaP cells treated with DMSO or 100 nM R1881. RNase H treatment as indicated. Shown is percent IP/Input as mean ± SEM for at least three replicates. RNase H treated IPs were performed once. D) Representative images of nuclei with *EWSR1* alleles intact (No break) or split (Break). White arrows indicate a break apart allele. E) Quantification of percent nuclei with a break. VCaP cells were treated with indicated doses of R1881 for 24 hours. TNFα treatment was for 48 hours. p values (*p < 0.05, **p < 0.001, ****p < 0.0001) were calculated by chi-square (N=85, 30, 67, 27, 25, 36, 37). F) Immunoblot of flEWS, ntEWS, and tubulin (TUB) in designated conditions. See also Figure S5.

### Androgen signaling promotes *EWSR1* breakpoint formation via R-loops

The *ntEWS* PAS is in close genomic proximity to the sequence in *EWSR1* that recurrently rearranges with *FLI1* in Ewing’s sarcoma, the breakpoint hotspot. From this observation, we developed the hypothesis that in addition to regulating *ntEWS* expression, AR could regulate *EWSR1* breakpoint formation. In fact, plotting the AR ChIP-seq dataset from VCaP cells shows AR binding flanks the breakpoint hotspot (Figure 5A). This was particularly curious since AR binding to *TMPRSS2* and *ERG* introns promotes formation of the *TMPRSS2/ERG* gene rearrangement in prostate cancer (Lin et al., 2009). Further, another nuclear hormone receptor, Estrogen Receptor, promotes breast cancer associated translocations by stimulating R-loop formation at target genes (Stork et al., 2016). Therefore, we investigated the importance of R-loops in *EWSR1* breakpoint formation as these can be a hotspot for DNA damage (Hamperl and Cimprich, 2014). Analysis of R-loops by DNA:RNA immunoprecipitation followed by next-generation sequencing (DRIP-seq) in a cell line of embryonic carcinoma of the testis (NTERA2) (Sanz and Chedin, 2019) showed that the same sequence in *EWSR1* that harbors the breakpoint hotspot forms an R-loop (Figure 5A). The breakpoint R-loop is resolved by treatment with Rnase H, which degrades the R-loop-associated RNA. To determine if androgen signaling impacts R-loop abundance genome wide, R-loop abundance was measured in VCaP cells treated with DMSO or 100 nM R1881 for 24 hours by dot blot using the S9.6 DNA:RNA hybrid antibody (Figure 5B). Treatment with R1881 caused an increase in R-loops genome wide and this signal was diminished by RNase H (Figure 5B). DRIP-qPCR showed an R1881-dependent increase in R-loops at the *EWSR1* locus, but an R1881 independent R-loop at the *CALM3* gene, which is a known R-loop hotspot (Figure 5C). Therefore, androgen signaling can promote increased R-loop formation at the *EWSR1* gene.

To test the role of androgen signaling in *EWSR1* breakpoint formation, a break apart fluorescence *in situ* hybridization assay (FISH) was used. Fluorescent probes flank the *EWSR1* breakpoint hotspot and create a merged red and green signal (yellow) when the *EWSR1* gene is intact and split signal when a break has occurred (Figure 5D). Treatment of VCaP cells with R1881 caused increased break formation in a dose dependent manner (Figure 5E), with a striking 80% of cells showing *EWSR1* breakage at the supraphysiological dose of 100 nM. Because high levels of androgen can cause DNA damage (Chatterjee et al., 2019), we asked if lower levels of androgen signaling could also promote high frequency breakage of *EWSR1*. Mani *et al.* reports that inflammation induced oxidative stress mediated by TNFα combined with androgen signaling promotes formation of the TMPRSS2/ERG fusion in prostate cells (Mani et al., 2016). We found that VCaP cells treated with TNFα alone did not show significantly increased *EWSR1* breakpoint frequency, however the combination of low dose (1 nM) R1881 with TNFα increased breakpoint frequency to near that of cells treated with high dose (100 nM) R1881 (Figure 5E). These data suggest that androgen signaling promotes *EWSR1* breaks alone at high doses or at low doses in collaboration with inflammation induced stress. To test the importance of R-loops in *EWSR1* breakpoint formation, RNase H was expressed in VCaP cells, treated with DMSO or 100 nM R1881 (Figure S5A), and these cells were assayed by break apart FISH. RNase H significantly abrogated breakpoint formation in cells with 100 nM R1881 (Figure 5E). This suggests that R-loops are essential for androgen-mediated *EWSR1* breakpoint formation.

Transcripts commonly terminate through the formation of transcription dependent R-loops (Sanz et al., 2016). RNase H expression in VCaP cells abolished expression of ntEWS and decreased expression of flEWS (Figure 5F). This is consistent with both 5’ and 3’ R-loops in the EWSR1 gene (Supplemental Figure 5c). These data suggest that R-loops can regulate both flEWS and ntEWS expression and can promote break-point formation.

## Discussion

We have shown in the prostate cancer setting that AR binds to intron 5 of *EWSR1* to directly upregulate a previously uncharacterized isoform that we have termed ntEWS. Our data indicate that ntEWS is localized to the cytoplasm and can strongly promote phenotypes associated with cancer such as cell migration and clonogenic growth. Further, AR signaling promoted increased R-loop formation in the EWSR1 gene and drove chromosomal breakage at high frequency at the same genomic locus that is rearranged in Ewing sarcoma.

Alternative RNA processing events can give rise to tissue specific gene isoforms (Lianoglou et al., 2013), and the misexpression of such isoforms is common in cancer (Simpson et al., 2005). Specifically, mRNA shortening through alternative polyadenylation is common in cancer (Mayr and Bartel, 2009). Yet exactly how RNA processing becomes dysregulated, what factors dictate this process in *cis* and in *trans*, and the downstream cellular consequences remain unclear. While 43.5% of AR binding sites are intronic (Wilson et al., 2016), the exact role of AR-bound introns is under-examined. In the context of *EWSR1*, our data indicate that intronic AR binding stimulates early termination. RNA processing is inherently coupled to transcription, and many transcription factors, including AR, interact with RNA processing factors. For example, the RNA processing factors NONO and SFPQ interact with AR to increase transcriptional output (Dong et al., 2007; Ishitani et al., 2003; Kuwahara et al., 2006). Future work is needed to test if AR can alter RNA processing via these or other factors.

We found that *ntEWS* expression is normally high in the testes. Because this organ is androgen regulated, this may explain why *ntEWS* expression is under control of AR. Mouse knockout models for both *EWSR1* and *AR* are defective in spermatogenesis due to meiotic arrest and increased apoptosis (De Gendt et al., 2004; Li et al., 2007), indicating a potential relationship between these genes in the testes. It is important to note that the Cre-Flox targeting strategy used in these studies to delete *EWSR1*, did eliminate essential exons for *ntEWS* (Li et al., 2007). Our data indicate that ntEWS functions in the cytoplasm, can bind cytoskeletal proteins and can form the higher order structure associated with phase separation. Interestingly, in full-length EWS, RNA can titrate the formation of higher order structures through binding the EWS RNA binding domain (Maharana et al., 2018). Since ntEWS lacks this RNA binding domain, higher-order structures formed by ntEWS would not be regulated by this mechanism.

Unlike many other gene fusions, we do not have a mechanistic explanation for why *EWSR1/FLI1* forms. Ewing’s sarcoma preferentially affects adolescent males with peak incidence at 15 years of age (Grunewald et al., 2018). This affected population is typically undergoing puberty, a period of peak androgen signaling. Our data suggest that this increase in androgen signaling, possibly combined with inflammatory signaling could promote the chromosomal breaks necessary for *EWSR1/FLI1* formation. The origins of the most common fusion in prostate cancer, *TMPRSS2/ERG* is accredited to intronic AR binding in combination with DNA damage or inflammatory signaling (Lin et al., 2009; Mani et al., 2016). Our current study suggests a similar mechanism could lead to *EWSR1* breakpoint formation followed by *EWSR1/FLI1* fusion. Since the *EWSR1* breakpoint hotspot is so close to the intronic PAS, we speculate that alternative polyadenylation stress may be related to breakpoint formation. Although we were able to generate *EWSR1* breakpoints at high frequencies in a prostate cancer cell line, *EWSR1* rearrangements are not common in prostate cancer. A recent study did find a *EWSR1/FEV* fusion in a prostate tumor from a single patient, but it is unclear whether this tumor should be categorized as prostate adenocarcinoma or Ewing’s sarcoma (Febres-Aldana et al., 2020). Our data indicate that high levels of AR in the cell are necessary for AR binding and function within the *EWSR1* gene. It is unclear if the cell of origin for Ewing sarcoma would express AR. While mesenchymal stem cells are a likely Ewing’s sarcoma cell of origin, the exact subtype of these cells that are targets and their state when *EWSR1/FLI1* forms is unknown. Future studies that stimulate both AR and inflammation in a mesenchymal cell setting are needed to test this potential mechanism for gene rearrangement.

## Methods

### RNA extraction and quantitative real time PCR

RNA was extracted using the RNAeasy kit (Qiagen) and DNase treated with the RNase-Free DNase kit (Qiagen) following manufacturer’s protocols. 1μg of total RNA was reverse transcribed using isoform specific primers (Table 1). RNA was measured by qRTPCR using standard curves as previously described (Hollenhorst et al., 2011). Expression was normalized to 18S and reported as three biological replicates each represented by the average of two technical replicates.

### Cell lines

VCaP, LNCaP, and PC3 cells were obtained from ATCC and passaged less than 20 times. All lines were cultured as according to manufacturer guidelines. In androgen depletion experiments, VCaPs and LNCaPs were androgen starved using phenol red free media supplemented with charcoal stripped FBS (Gibco) for 48hr prior to R1881 (Sigma) exposure.

### Viral transductions and transient transfections

Overexpression constructs were expressed in PC3s by retrovirus and have N-terminal 2xHA tags. Full length EWS was described previously (Kedage et al., 2016), the N-terminal EWS isoform was cloned from LNCaP total RNA. The C-terminal EWS isoform was cloned from full length. AR was cloned from pCMV-FLAG-hAR(Bai et al., 2005) (a gift from Elizabeth Wilson; Addgene plasmid #89080; http://net.net/addgene89080; RRID:Addgene_89080). See Table 1 for cloning primers. The RNase H expression construct (ppyCAG_RNaseH1_WT(Chen et al., 2017) was a gift from Xiang-Dong Fu; Addgene plasmid #111906; http://n2t.net/addgene:111906; RRID Addgene_111906) was expressed in VCaP cells via transient transfection using TransIT 20-20 (Mirus). Cells were split two days post transfection for downstream experiments.

### Clonogenic growth, cell migration, and MTT assays

Transwell migration assays were preformed as previously described (Hollenhorst et al., 2011). Briefly, PC3s were plated into transwells (8 μM pore size, BD Bioscience) at a density of 500,000 cells/well in serum free media and allowed 48 hr to migrate towards serum containing media. Migrated cells were then fixed, stained, and quantified; values are the mean and SEM of three biological replicates with two technical replicates each. Clonogenic growth was performed as previously described (Kedage et al., 2016). 1,000 PC3 cells were seeded into a well of a 6-well plate and allowed to grow for 10 days. Colonies were fixed with 10% formalin, stained with 0.5% crystal violet (Sigma) in 25% methanol, and counted using Genesys software (Syngene). Reported colonies are the mean of three biological replicates with two technical replicates each. Cell proliferation was measured using the MTT (Calbiochem) assay as previously described (Plotnik et al., 2014). Briefly, 800 PC3 cells were seeded in a 96-well plate. Readings were taken 24 hr after plating with time points continuing for 4 days. MTT reagent (5mg/ml in PBS) was added to cells and incubated for four hours after which the MTT containing media was removed and DMSO added. After gentle agitation, absorbance (600nm) was measured using a microplate reader (ELx8200, BioTek Instruments). Cell proliferation was reported as three biological replicates each with five technical replicates.

### Precipitation assay

Protein precipitation via b-isox was performed as shown previously (Boulay et al., 2017). Briefly, cell lysates were divided into four tubes and treated with 0, 10, 30, or 100 μM b-isox (Sigma). Lysates were incubated at 4C with rotation for one hr. Protein was precipitated by centrifugation and washed twice and protein aggregation was monitored by immunoblot.

### ChIP

ChIP of indicated proteins was previously described (Hollenhorst et al., 2007) but after two days of androgen depletion and subsequent treatment with 10 nM R1881 or DMSO. Antibodies used for immunoblot were also used for ChIP. Briefly, crosslinking was carried out using 1% formaldehyde and quenched with glycine. Cells were lysed with ChIP cell lysis buffer (50 mM Hepes-KOH pH 8, 1 mM EDTA, 0.5 mM EGTA, 140 mM NaCl, 10% Glycerol, 0.5% NP-40, 0.25% Triton X-100) supplemented with protease inhibitors (Sigma) and nuclei were isolated and washed with ChIP wash buffer (10 mM Tris-HCL pH 8, 1 mM EDTA, 0.5 mM EGTA, 200 mM NaCl) supplemented with protease inhibitors (Sigma). Nuclei were then sonicated using a Biorupter Pico for 30 sec on, 30 sec off for 4 rounds. Nuclear extract was added to dynabead-antibody conjugates and rotated for 4 hours at 4 degrees. Bead complexes were washed with IP wash buffer (20 mM Tris pH 7.9, 0.25% NP-40, 0.05% SDS, 2 mM EDTA, 250 mM NaCl) four times. Protein and RNA were degraded by Proteinase K (Sigma) and RNase A (5prime), respectively. DNA was purified by phenol chloroform extraction and QiaQuick PCR Purification Kit (Qiagen).

### ChIP-seq analysis

Raw ChIP-seq fastq files for AR from patient tumors and matched adjacent normal tissue (GSE70079), for TFs in VCaPs (GSE56086), and for TFs in LNCaPs (GSE83860) were downloaded from the SRA using SRA toolkit (http://ncbi.github.io/sra-tools/). When applicable, sequencing replicates and biological replicates were concatenated. Reads were aligned using Bowtie2 (Langmead and Salzberg, 2012) and peaks were called using the default settings with MACS2 (Zhang et al., 2008) after removing PCR duplicates with samtools (Li et al., 2009). Motif analysis was performed with the AR motif (http://homer.ucsd.edu/homer/motif/HomerMotifDB/homerResults/motif6.info.html), AR half-site (http://homer.ucsd.edu/homer/motif/HomerMotifDB/homerResults/motif7.info.html), and FOXA1:AR motif (http://homer.ucsd.edu/homer/motif/HomerMotifDB/homerResults/motif5.info.html) using the FIMO tool on MEME-suite (Grant et al., 2011).

### Co-immunoprecipation assay for mass spectrometry

All co-immunoprecipitations were preformed using cytoplasmic extracts. PC3 cells were washed with PBS, harvested in cell lysis buffer (50 mM Hepes-KOH pH 8, 1 mM EDTA, 0.5 mM EGTA, 140 mM NaCl, 10% Glycerol, 0.5% NP-40, 0.25% Triton X-100) supplemented with protease inhibitors (Sigma) and rotated for 10 min at 4C. The cytoplasmic fraction was separated from nuclei by centrifugation at 700 rcf for 5 min. A Bradford assay determined total protein concentration. For downstream mass spectrometry analysis, at least 4 mg of protein was added to beads conjugated to anti-HA antibodies. Cytoplasmic extracts were then incubated with bead-antibody conjugates for 24 hrs at 4C with rotation and washed four times with PBS and sent to the Indiana University School of Medicine Proteomics Core for identification of interacting partners. Interactions that scored a spectrum count of log2(ntEWS/EWS(1-355aa)) < −1.5 in both replicates were called as EWS (1-355aa) enriched; and interactions that scored a spectrum count of log2(ntEWS/EWS(1-355aa)) > 1.5 were called as ntEWS enriched. Proteins that had a spectrum count of −1.5 < log2(ntEWS/EWS(1-355aa)) > 1.5 were considered to be shared interacting partners.

### Dot Blot

Nitrocellulose membrane was soaked in 6x SSC buffer (diluted in ddH_2_O from 20x; 3 M NaCl, 300mM trisodium citrate, pH 7) for 30 min to 1 hr then assembled in a dot blot apparatus (BioRad). After washing with 100 μl TE buffer, 2 μg and 1 μg of digested genomic DNA in 100 μl of TE buffer was added to appropriate wells and vacuum filtered. Wells were washed twice with 2x SSC buffer (diluted in ddH_2_O from 20x). The membrane was washed for 30 sec with 2x SSC buffer, air dried for 30 min, crosslinked using 0.12J/m^2^ UV, then immunoblotted with S9.6 antibody (Abcam) at 1:1000 dilution.

### DRIP-qPCR

DRIP-qPCR was done as previously described (Sanz and Chedin, 2019) with the following changes. Genomic DNA was extracted from ∼3 million VCaP cells, immunoprecipitated with 8 μl of S9.6 antibody and 100ul anti-mouse Dynabeads (Invitrogen), and DNA cleaned up using AMPure beads (Beckman Coulter Inc.). Enrichment is reported as percent IP/Input.

### Break-apart FISH

The *EWSR1* break-apart FISH probes were made by Lecia Biosystems. Break-apart FISH was performed as previously described (Mittal, 2015) with the exception of incubating cells with 2x SSC for 5 min at 37C instead of 30 min. *EWSR1* is on chromosome 22; VCaP cells have three copies of this chromosome, thus signal from all alleles was required for scoring. Breaks were scored by split green-red signal by at least one signal diameter as shown previously (Mittal, 2015).

## Acknowledgements

Research reported in this publication was supported by the National Cancer Institute of the National Institutes of Health under Award Number R01CA204121 (P.C.H.), and with support from the Indiana Clinical and Translational Sciences Institute funded, in part by award number UL1TR002529 from the National Institutes of Health, National Center for Advancing Translational Sciences, Clinical and Translational Sciences Award (T.R.N.). Funding was also provided by the Doane and Eunice Dahl Wright Fellowship from the Indiana University School of Medicine (T.R.N.). Mass spectrometry was provided by the Indiana University School of Medicine Proteomics Core Facility. Microscopy assistance was provided by the Indiana University Light Microscopy Imaging Center. The authors would like to thank Shruthi Sriramkumar for advice on the dot blot.

## References

Altmeyer, M., Neelsen, K.J., Teloni, F., Pozdnyakova, I., Pellegrino, S., Grofte, M., Rask, M.D., Streicher, W., Jungmichel, S., Nielsen, M.L., et al. (2015). Liquid demixing of intrinsically disordered proteins is seeded by poly(ADP-ribose). Nat Commun 6, 8088.

Andersson, M.K., Stahlberg, A., Arvidsson, Y., Olofsson, A., Semb, H., Stenman, G., Nilsson, O., and Aman, P. (2008). The multifunctional FUS, EWS and TAF15 proto-oncoproteins show cell type-specific expression patterns and involvement in cell spreading and stress response. BMC Cell Biol 9, 37.

Azuma, M., Embree, L.J., Sabaawy, H., and Hickstein, D.D. (2007). Ewing sarcoma protein ewsr1 maintains mitotic integrity and proneural cell survival in the zebrafish embryo. PLoS One 2, e979.

Bai, S., He, B., and Wilson, E.M. (2005). Melanoma antigen gene protein MAGE-11 regulates androgen receptor function by modulating the interdomain interaction. Mol Cell Biol 25, 1238–1257.

Boulay, G., Sandoval, G.J., Riggi, N., Iyer, S., Buisson, R., Naigles, B., Awad, M.E., Rengarajan, S., Volorio, A., McBride, M.J., et al. (2017). Cancer-Specific Retargeting of BAF Complexes by a Prion-like Domain. Cell 171, 163–178 e119.

Chatterjee, P., Schweizer, M.T., Lucas, J.M., Coleman, I., Nyquist, M.D., Frank, S.B., Tharakan, R., Mostaghel, E., Luo, J., Pritchard, C.C., et al. (2019). Supraphysiological androgens suppress prostate cancer growth through androgen receptor-mediated DNA damage. J Clin Invest 130, 4245–4260.

Chen, L., Chen, J.Y., Zhang, X., Gu, Y., Xiao, R., Shao, C., Tang, P., Qian, H., Luo, D., Li, H., et al. (2017). R-ChIP Using Inactive RNase H Reveals Dynamic Coupling of R-loops with Transcriptional Pausing at Gene Promoters. Mol Cell 68, 745–757 e745.

Couthouis, J., Hart, M.P., Erion, R., King, O.D., Diaz, Z., Nakaya, T., Ibrahim, F., Kim, H.J., Mojsilovic-Petrovic, J., Panossian, S., et al. (2012). Evaluating the role of the FUS/TLS-related gene EWSR1 in amyotrophic lateral sclerosis. Hum Mol Genet 21, 2899–2911.

De Gendt, K., Swinnen, J.V., Saunders, P.T., Schoonjans, L., Dewerchin, M., Devos, A., Tan, K., Atanassova, N., Claessens, F., Lecureuil, C., et al. (2004). A Sertoli cell-selective knockout of the androgen receptor causes spermatogenic arrest in meiosis. Proc Natl Acad Sci U S A 101, 1327–1332.

Deloulme, J.C., Prichard, L., Delattre, O., and Storm, D.R. (1997). The prooncoprotein EWS binds calmodulin and is phosphorylated by protein kinase C through an IQ domain. J Biol Chem 272, 27369–27377.

Diao, Y., Fang, R., Li, B., Meng, Z., Yu, J., Qiu, Y., Lin, K.C., Huang, H., Liu, T., Marina, R.J., et al. (2017). A tiling-deletion-based genetic screen for cis-regulatory element identification in mammalian cells. Nat Methods 14, 629–635.

Dong, X., Sweet, J., Challis, J.R., Brown, T., and Lye, S.J. (2007). Transcriptional activity of androgen receptor is modulated by two RNA splicing factors, PSF and p54nrb. Mol Cell Biol 27, 4863–4875.

Dosztanyi, Z. (2018). Prediction of protein disorder based on IUPred. Protein Sci 27, 331–340.

Dutertre, M., Sanchez, G., De Cian, M.C., Barbier, J., Dardenne, E., Gratadou, L., Dujardin, G., Le Jossic-Corcos, C., Corcos, L., and Auboeuf, D. (2010). Cotranscriptional exon skipping in the genotoxic stress response. Nat Struct Mol Biol 17, 1358–1366.

Febres-Aldana, C.A., Krishnamurthy, K., Delgado, R., Kochiyil, J., Poppiti, R., and Medina, A.M. (2020). Prostatic carcinoma with neuroendocrine differentiation harboring the EWSR1-FEV fusion transcript in a man with the WRN G327X germline mutation: A new variant of prostatic carcinoma or a member of the Ewing sarcoma family of tumors? Pathol Res Pract 216, 152758.

Felsch, J.S., Lane, W.S., and Peralta, E.G. (1999). Tyrosine kinase Pyk2 mediates G-protein-coupled receptor regulation of the Ewing sarcoma RNA-binding protein EWS. Curr Biol 9, 485–488.

Gasperini, M., Findlay, G.M., McKenna, A., Milbank, J.H., Lee, C., Zhang, M.D., Cusanovich, D.A., and Shendure, J. (2017). CRISPR/Cas9-Mediated Scanning for Regulatory Elements Required for HPRT1 Expression via Thousands of Large, Programmed Genomic Deletions. Am J Hum Genet 101, 192–205.

Goldman, M.C. B.; Kamath, A.; Brooks, A.; Zhu, J.; Haussler, D. (2018). The UCSC Xena Platform for cancer genomics data visualization and interpretation. bioRxiv.

Gorthi, A., Romero, J.C., Loranc, E., Cao, L., Lawrence, L.A., Goodale, E., Iniguez, A.B., Bernard, X., Masamsetti, V.P., Roston, S., et al. (2018). EWS-FLI1 increases transcription to cause R-loops and block BRCA1 repair in Ewing sarcoma. Nature 555, 387–391.

Grant, C.E., Bailey, T.L., and Noble, W.S. (2011). FIMO: scanning for occurrences of a given motif. Bioinformatics 27, 1017–1018.

Grunewald, T.G.P., Cidre-Aranaz, F., Surdez, D., Tomazou, E.M., de Alava, E., Kovar, H., Sorensen, P.H., Delattre, O., and Dirksen, U. (2018). Ewing sarcoma. Nat Rev Dis Primers 4, 5.

Haeussler, M., Zweig, A.S., Tyner, C., Speir, M.L., Rosenbloom, K.R., Raney, B.J., Lee, C.M., Lee, B.T., Hinrichs, A.S., Gonzalez, J.N., et al. (2019). The UCSC Genome Browser database: 2019 update. Nucleic Acids Res 47, D853–D858.

Haffner, M.C., Aryee, M.J., Toubaji, A., Esopi, D.M., Albadine, R., Gurel, B., Isaacs, W.B., Bova, G.S., Liu, W., Xu, J., et al. (2010). Androgen-induced TOP2B-mediated double-strand breaks and prostate cancer gene rearrangements. Nat Genet 42, 668–675.

Hamperl, S., and Cimprich, K.A. (2014). The contribution of co-transcriptional RNA:DNA hybrid structures to DNA damage and genome instability. DNA Repair (Amst) 19, 84–94.

Hollenhorst, P.C., Paul, L., Ferris, M.W., and Graves, B.J. (2011). The ETS gene ETV4 is required for anchorage-independent growth and a cell proliferation gene expression program in PC3 prostate cells. Genes Cancer 1, 1044–1052.

Hollenhorst, P.C., Shah, A.A., Hopkins, C., and Graves, B.J. (2007). Genome-wide analyses reveal properties of redundant and specific promoter occupancy within the ETS gene family. Genes Dev 21, 1882–1894.

Huang, C.K., Luo, J., Lee, S.O., and Chang, C. (2014). Concise review: androgen receptor differential roles in stem/progenitor cells including prostate, embryonic, stromal, and hematopoietic lineages. Stem Cells 32, 2299–2308.

Ishitani, K., Yoshida, T., Kitagawa, H., Ohta, H., Nozawa, S., and Kato, S. (2003). p54nrb acts as a transcriptional coactivator for activation function 1 of the human androgen receptor. Biochem Biophys Res Commun 306, 660–665.

Kapeli, K., Martinez, F.J., and Yeo, G.W. (2017). Genetic mutations in RNA-binding proteins and their roles in ALS. Hum Genet 136, 1193–1214.

Kato, M., Han, T.W., Xie, S., Shi, K., Du, X., Wu, L.C., Mirzaei, H., Goldsmith, E.J., Longgood, J., Pei, J., et al. (2012). Cell-free formation of RNA granules: low complexity sequence domains form dynamic fibers within hydrogels. Cell 149, 753–767.

Kedage, V., Selvaraj, N., Nicholas, T.R., Budka, J.A., Plotnik, J.P., Jerde, T.J., and Hollenhorst, P.C. (2016). An Interaction with Ewing’s Sarcoma Breakpoint Protein EWS Defines a Specific Oncogenic Mechanism of ETS Factors Rearranged in Prostate Cancer. Cell Rep 17, 1289–1301.

Kelley, L.A., Mezulis, S., Yates, C.M., Wass, M.N., and Sternberg, M.J. (2015). The Phyre2 web portal for protein modeling, prediction and analysis. Nat Protoc 10, 845–858.

Kuwahara, S., Ikei, A., Taguchi, Y., Tabuchi, Y., Fujimoto, N., Obinata, M., Uesugi, S., and Kurihara, Y. (2006). PSPC1, NONO, and SFPQ are expressed in mouse Sertoli cells and may function as coregulators of androgen receptor-mediated transcription. Biol Reprod 75, 352–359.

Langmead, B., and Salzberg, S.L. (2012). Fast gapped-read alignment with Bowtie 2. Nat Methods 9, 357–359.

Li, H., Handsaker, B., Wysoker, A., Fennell, T., Ruan, J., Homer, N., Marth, G., Abecasis, G., Durbin, R., and Genome Project Data Processing, S. (2009). The Sequence Alignment/Map format and SAMtools. Bioinformatics 25, 2078–2079.

Li, H., Watford, W., Li, C., Parmelee, A., Bryant, M.A., Deng, C., O’Shea, J., and Lee, S.B. (2007). Ewing sarcoma gene EWS is essential for meiosis and B lymphocyte development. J Clin Invest 117, 1314–1323.

Lianoglou, S., Garg, V., Yang, J.L., Leslie, C.S., and Mayr, C. (2013). Ubiquitously transcribed genes use alternative polyadenylation to achieve tissue-specific expression. Genes Dev 27, 2380–2396.

Lin, C., Yang, L., Tanasa, B., Hutt, K., Ju, B.G., Ohgi, K., Zhang, J., Rose, D.W., Fu, X.D., Glass, C.K., et al. (2009). Nuclear receptor-induced chromosomal proximity and DNA breaks underlie specific translocations in cancer. Cell 139, 1069–1083.

Lonergan, P.E., and Tindall, D.J. (2011). Androgen receptor signaling in prostate cancer development and progression. J Carcinog 10, 20.

Maharana, S., Wang, J., Papadopoulos, D.K., Richter, D., Pozniakovsky, A., Poser, I., Bickle, M., Rizk, S., Guillen-Boixet, J., Franzmann, T.M., et al. (2018). RNA buffers the phase separation behavior of prion-like RNA binding proteins. Science 360, 918–921.

Malinen, M., Niskanen, E.A., Kaikkonen, M.U., and Palvimo, J.J. (2017). Crosstalk between androgen and pro-inflammatory signaling remodels androgen receptor and NF-kappaB cistrome to reprogram the prostate cancer cell transcriptome. Nucleic Acids Res 45, 619–630.

Mani, R.S., Amin, M.A., Li, X., Kalyana-Sundaram, S., Veeneman, B.A., Wang, L., Ghosh, A., Aslam, A., Ramanand, S.G., Rabquer, B.J., et al. (2016). Inflammation-Induced Oxidative Stress Mediates Gene Fusion Formation in Prostate Cancer. Cell Rep 17, 2620–2631.

Mayr, C., and Bartel, D.P. (2009). Widespread shortening of 3’UTRs by alternative cleavage and polyadenylation activates oncogenes in cancer cells. Cell 138, 673–684.

Mittal, N., Kunz, C., Gypas, F., Kishore, S., Martin, G., Wenzel, F., van Nimwegen, E., Schar, R., Zavolan, M. (2015). Ewing sarcoma breakpoint region 1 prevents transcription-associated genome instability. bioRxiv.

Murashima, A., Kishigami, S., Thomson, A., and Yamada, G. (2015). Androgens and mammalian male reproductive tract development. Biochim Biophys Acta 1849, 163–170.

Nyquist, K.B., Thorsen, J., Zeller, B., Haaland, A., Troen, G., Heim, S., and Micci, F. (2011). Identification of the TAF15-ZNF384 fusion gene in two new cases of acute lymphoblastic leukemia with a t(12;17)(p13;q12). Cancer Genet 204, 147–152.

Pahlich, S., Quero, L., Roschitzki, B., Leemann-Zakaryan, R.P., and Gehring, H. (2009). Analysis of Ewing sarcoma (EWS)-binding proteins: interaction with hnRNP M, U, and RNA-helicases p68/72 within protein-RNA complexes. J Proteome Res 8, 4455–4465.

Paronetto, M.P., Minana, B., and Valcarcel, J. (2011). The Ewing sarcoma protein regulates DNA damage-induced alternative splicing. Mol Cell 43, 353–368.

Patel, A., Lee, H.O., Jawerth, L., Maharana, S., Jahnel, M., Hein, M.Y., Stoynov, S., Mahamid, J., Saha, S., Franzmann, T.M., et al. (2015). A Liquid-to-Solid Phase Transition of the ALS Protein FUS Accelerated by Disease Mutation. Cell 162, 1066–1077.

Plotnik, J.P., Budka, J.A., Ferris, M.W., and Hollenhorst, P.C. (2014). ETS1 is a genome-wide effector of RAS/ERK signaling in epithelial cells. Nucleic Acids Res 42, 11928–11940.

Pomerantz, M.M., Li, F., Takeda, D.Y., Lenci, R., Chonkar, A., Chabot, M., Cejas, P., Vazquez, F., Cook, J., Shivdasani, R.A., et al. (2015). The androgen receptor cistrome is extensively reprogrammed in human prostate tumorigenesis. Nat Genet 47, 1346–1351.

Qamar, S., Wang, G., Randle, S.J., Ruggeri, F.S., Varela, J.A., Lin, J.Q., Phillips, E.C., Miyashita, A., Williams, D., Strohl, F., et al. (2018). FUS Phase Separation Is Modulated by a Molecular Chaperone and Methylation of Arginine Cation-pi Interactions. Cell 173, 720–734 e715.

Riggi, N., Cironi, L., Suva, M.L., and Stamenkovic, I. (2007). Sarcomas: genetics, signalling, and cellular origins. Part 1: The fellowship of TET. J Pathol 213, 4–20.

Rossow, K.L., and Janknecht, R. (2001). The Ewing’s sarcoma gene product functions as a transcriptional activator. Cancer Res 61, 2690–2695.

Sanz, L.A., and Chedin, F. (2019). High-resolution, strand-specific R-loop mapping via S9.6-based DNA-RNA immunoprecipitation and high-throughput sequencing. Nat Protoc 14, 1734–1755.

Sanz, L.A., Hartono, S.R., Lim, Y.W., Steyaert, S., Rajpurkar, A., Ginno, P.A., Xu, X., and Chedin, F. (2016). Prevalent, Dynamic, and Conserved R-Loop Structures Associate with Specific Epigenomic Signatures in Mammals. Mol Cell 63, 167–178.

Shaw, D.J., Morse, R., Todd, A.G., Eggleton, P., Lorson, C.L., and Young, P.J. (2009). Identification of a tripartite import signal in the Ewing Sarcoma protein (EWS). Biochem Biophys Res Commun 390, 1197–1201.

Simpson, A.J., Caballero, O.L., Jungbluth, A., Chen, Y.T., and Old, L.J. (2005). Cancer/testis antigens, gametogenesis and cancer. Nat Rev Cancer 5, 615–625.

Stork, C.T., Bocek, M., Crossley, M.P., Sollier, J., Sanz, L.A., Chedin, F., Swigut, T., and Cimprich, K.A. (2016). Co-transcriptional R-loops are the main cause of estrogen-induced DNA damage. Elife 5.

Tanikawa, C., Ueda, K., Suzuki, A., Iida, A., Nakamura, R., Atsuta, N., Tohnai, G., Sobue, G., Saichi, N., Momozawa, Y., et al. (2018). Citrullination of RGG Motifs in FET Proteins by PAD4 Regulates Protein Aggregation and ALS Susceptibility. Cell Rep 22, 1473–1483.

Tate, J.G., Bamford, S., Jubb, H.C., Sondka, Z., Beare, D.M., Bindal, N., Boutselakis, H., Cole, C.G., Creatore, C., Dawson, E., et al. (2019). COSMIC: the Catalogue Of Somatic Mutations In Cancer. Nucleic Acids Res 47, D941–D947.

Thomas, J.D., Polaski, J.T., Feng, Q., De Neef, E.J., Hoppe, E.R., McSharry, M.V., Pangallo, J., Gabel, A.M., Belleville, A.E., Watson, J., et al. (2020). RNA isoform screens uncover the essentiality and tumor-suppressor activity of ultraconserved poison exons. Nat Genet 52, 84–94.

Toropainen, S., Malinen, M., Kaikkonen, S., Rytinki, M., Jaaskelainen, T., Sahu, B., Janne, O.A., and Palvimo, J.J. (2015). SUMO ligase PIAS1 functions as a target gene selective androgen receptor coregulator on prostate cancer cell chromatin. Nucleic Acids Res 43, 848–861.

Wang, R., Zheng, D., Yehia, G., and Tian, B. (2018). A compendium of conserved cleavage and polyadenylation events in mammalian genes. Genome Res 28, 1427–1441.

Wilson, S., Qi, J., and Filipp, F.V. (2016). Refinement of the androgen response element based on ChIP-Seq in androgen-insensitive and androgen-responsive prostate cancer cell lines. Sci Rep 6, 32611.

Yamamoto-Shiraishi, Y., Higuchi, H., Yamamoto, S., Hirano, M., and Kuroiwa, A. (2014). Etv1 and Ewsr1 cooperatively regulate limb mesenchymal Fgf10 expression in response to apical ectodermal ridge-derived fibroblast growth factor signal. Dev Biol 394, 181–190.

Zhang, Y., Liu, T., Meyer, C.A., Eeckhoute, J., Johnson, D.S., Bernstein, B.E., Nusbaum, C., Myers, R.M., Brown, M., Li, W., et al. (2008). Model-based analysis of ChIP-Seq (MACS). Genome Biol 9, R137.

Zhou, Y., Zhou, B., Pache, L., Chang, M., Khodabakhshi, A.H., Tanaseichuk, O., Benner, C., and Chanda, S.K. (2019). Metascape provides a biologist-oriented resource for the analysis of systems-level datasets. Nat Commun 10, 1523.

