## Supplemental figures and tables for "Androgen signaling connects short isoform production to breakpoint formation at Ewing sarcoma breakpoint region 1 via an R-loop-dependent mechanism"

A

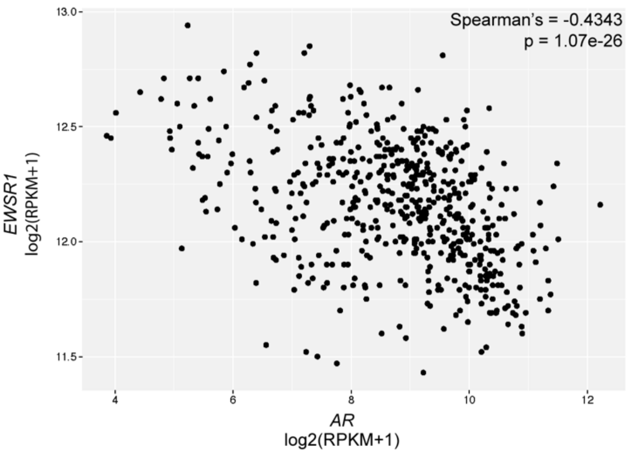


B

| | % of samples with expression | PAS signal | Mean RPM | Location |
|---|---|---|---|---|
| 1 | 4.7 | AAUAAA | 0.8 | Intron |
| 2 | 69.2 | AUUAAA | 3.0 | Intron |
| 3 | 2.8 | None | 1.4 | Intron |
| 4 | 1.9 | None | 1.8 | Coding exon |
| 5 | 77.6 | AUUAAA | 3.5 | 3' exon |
| 6 | 57.9 | None | 3.4 | 3' exon |
| 7 | 55.1 | AAUAAA | 3.0 | 3' exon |

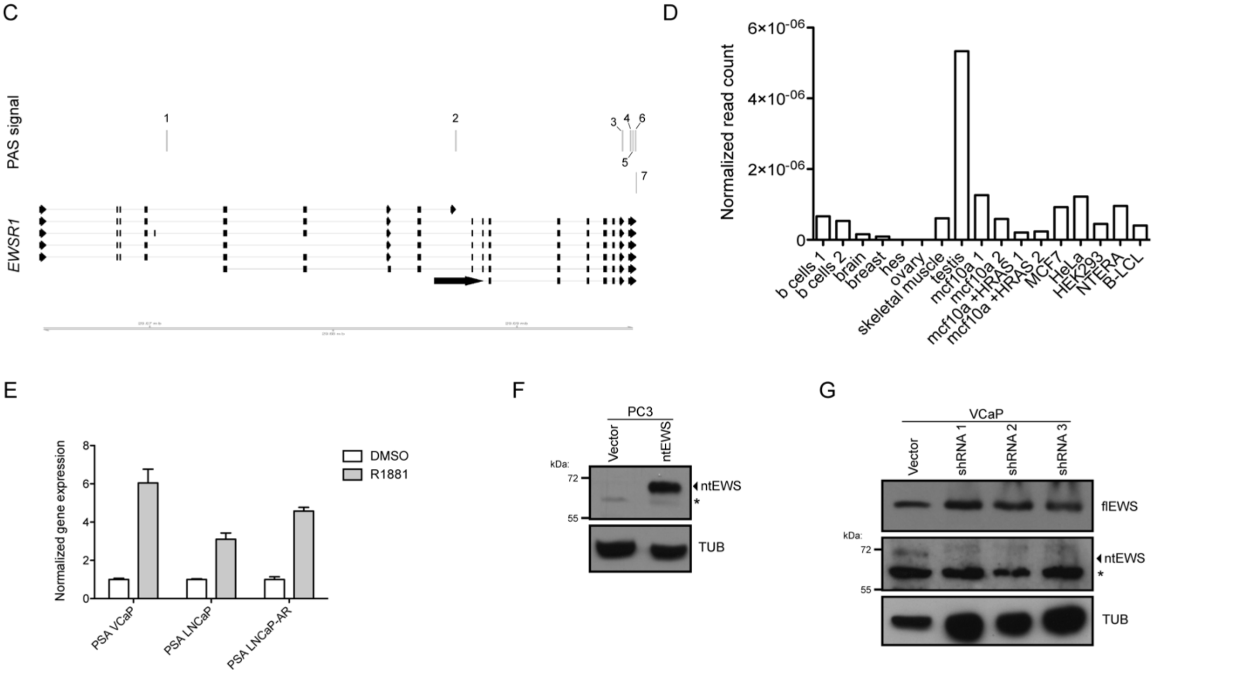


**Figure S1. Androgen signaling upregulates an intronic polyadenylated *EWSR1* isoform**

A) Gene expression correlation of *EWSR1* and *AR* in 550 prostate cancer patients from the PRAD data set. B) *EWSR1* polyadenylation site (PAS) information from PolyA_db v3.2. There are seven PAS for this gene and they are numbered starting from the 5' end of the gene. C) Diagram of PAS locations at *EWSR1* numbered according to table in A. D) Normalized read count for PAS #2, the PAS for *ntEWS*, from 3' sequencing data across various cell types. E) RNA levels of PSA in VCaP, LNCaP, and LNCaP-AR cells treated with DMSO or 10nM R1881 for 24 hours. Expression is normalized to 18S and relative to the DMSO condition. The mean ± SEM for three replicates is shown. F) Immuno blot of ntEWS in PC3 cells overexpressing HA-ntEWS. G) Immunoblot of ntEWS in VCaP cells overexpressing vector alone or shRNAs targeting the 3' UTR of ntEWS. Tubulin is used as a loading control for immunoblots.

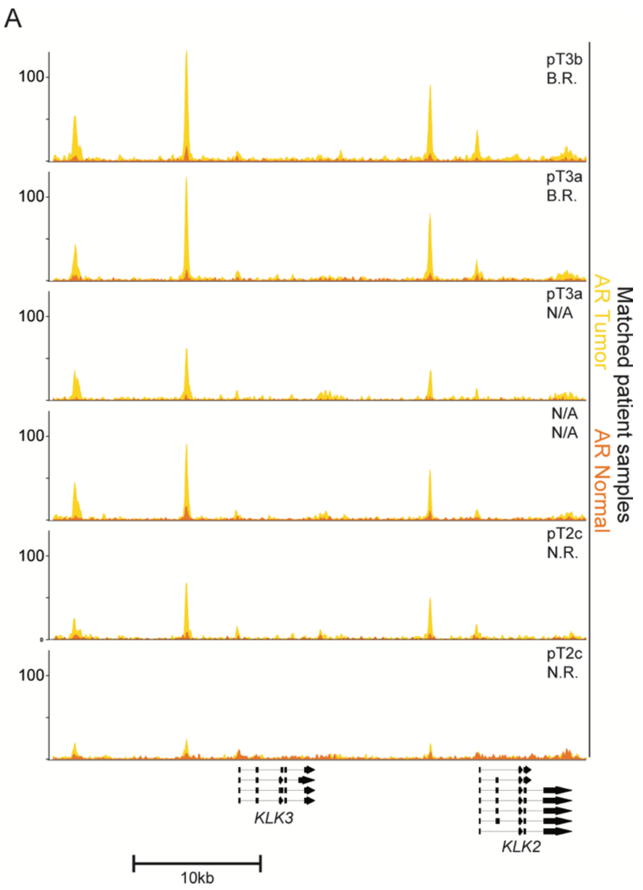


**Figure S2. AR binding to Intron 5 of *EWSR1* directly regulates ntEWS expression**

A) Gene tracks for AR binding in patient tumor and matched adjacent normal tissue at known AR enhancers. Order of tracks is consistent with Figure 2a.

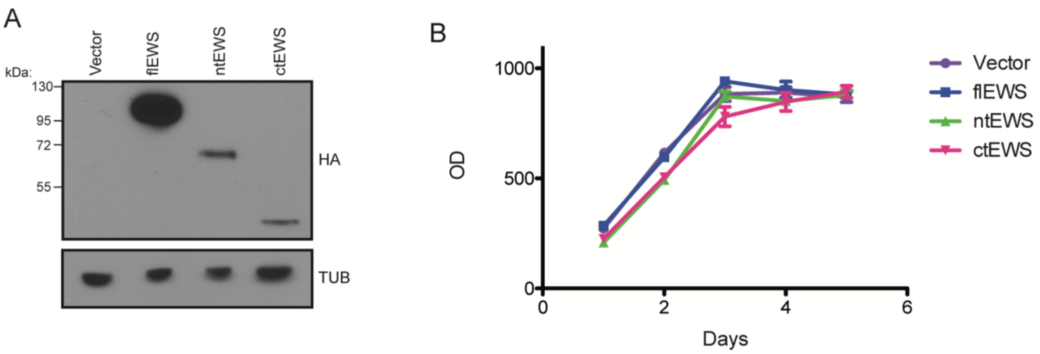


**Figure S3. ntEWS promotes phenotypes related to oncogenesis**

A) Immunoblot of 3xHA tagged EWS isoforms expressed in PC3 cells. Tubulin is used as a loading control. B) MTT proliferation assay of PC3 isoform-expressing lines.

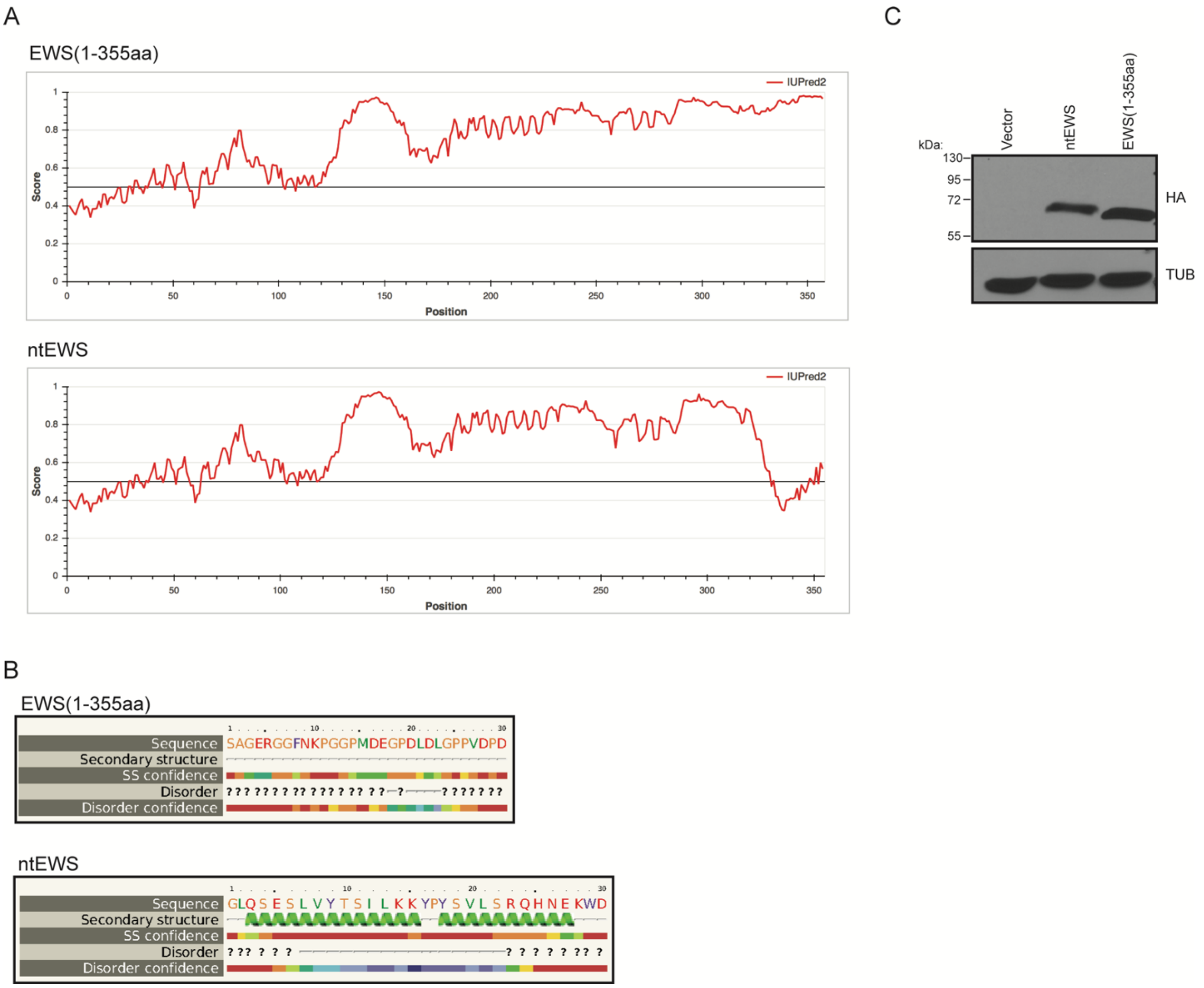


**Figure S4. The ntEWS alternative last exon encodes an alpha helical domain important for function**

A) IUPRED prediction of disorder of ntEWS (bottom) and EWS(1-355aa) (top). A score of 1 indicates intrinsic disorder. Scores below the line at 0.5 indicate structure. B) PHYRE2 analysis of the 30aa (324-354aa) encoded by the ALE of ntEWS (bottom) and 325-355aa of flEWS (top). C) Immunoblot of 3xHA tagged ntEWS and EWS (1-355aa) expressed in PC3 cells in addition to empty vector. Tubulin is used as a loading control.

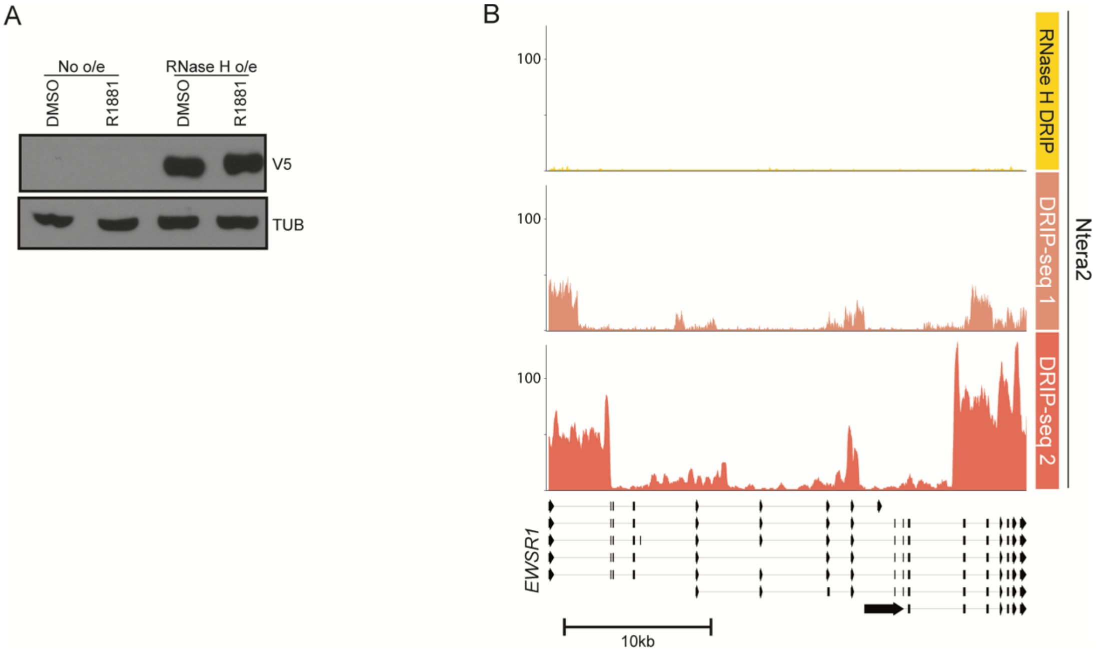


**Figure S5. Androgen signaling promotes *EWSR1* breakpoint formation via R-loops**
A) Immunoblot for V5-tagged RNase H in VCaP cells either treated with DMSO or 10nM R1881. Tubulin is used as a loading control. B) DRIP-seq tracks at *EWSR1* as labeled.

**Table S1: Primers used**

| | 5’ | 3’ |
|---|---|---|
| **Cloning primers** | | |
| HA-ntEWS | CTATGCATACCCATACGATGTTCCAGATTACGCTAAGGCGTCCACGGATTACAGTACCTAT | GTGACATTAATTAACTAGTCCCACTTTTCATTATGCTGCCG |
| HA-ctEWS | CTATGCATACCCATACGATGTTCCAGATTACGCTAAGGATGAAGGACCAGATCTTGAT | GTGACATTAATTAATTACTAGTAGGGCCGATCTCTGCG |
| HA-EWS (1-355aa) | CTATGCATACCCATACGATGTTCCAGATTACGCTAAGGCGTCCACGGATTACAGTACCTAT | GTGACATTAATTAATTACTAGTAGGGCCGATCTCTGCG |
| 3xHA | AGACTGCGGCCGCATGTATCCGTATGACGTCCCGGACTATGCATATCCGTATGACGTCCCGGACTATGCATACCCATACGATGTTC | |
| AR | AGACTGCGGCCGCATGGAAGTGCAGTTAGGGCTG | GTGACATTAATTAATCACTGGGTGTGGAAATAGAT |
| **RT-qPCR primers** | | |
| flEWS | TTATGGGCAGGAGTCTGGAGG | CTGGTCCTTCATCCATGGGTC |
| ntEWS 1 | GGAGGATTTTCCGGACCAGG | GTATACAAGGCTCTCACTTTG |
| ntEWS 2 | GGAGGATTTTCCGGACCAGG | CTAGTCCCACTTTTCATTATGC |
| PSA | GTGACCAAGTTCATGCTGTG | TTGGCCACGATGGTGTCCTTG |
| 18S | GGTGAAATTCTTGGACCGGC | GACTTTGGTTTCCCGGAAGC |
| **ChIP-qPCR primers** | | |
| Intron 5 site | GCGTTTACTGTGATGAATGGAGC | CTCCTGGGTAAGAATGCTAC |
| Intron 8 site | TGCATGCAACAGCTTGAAAT | GAGGGGAGAGGGAAATATGAA |
| XKRT (neg) | GGGATGGAGGTTTGCTCTTG | TGGACATGGTAGCGGGTAC |
| **gRNA primers** | | |
| Downstream FOXA1:AR site | CACCGAGCTTTGTAGCATTCTTACCC | AAACGGGTAAGAATGCTACAAAGCTC |
| Upstream FOXA1:AR site | CACCGATCCGGGAGAAGTGATCTGTT | AAACAACAGATCACTTCTCCCGGATC |
| **DRIP-qPCR primers** | | |
| CALM3 (Sanz and Chedin, 2019) | GAGGAATTGTGGCGTTGACT | AGAGTGGCCAAATGAGCAGT |
| EWS | CCTTGGTTAGTGCCTTGGAA | GTCGGAATGAACCTGAGGAA |

**Table S2: IP-MS identified interacting partners**

| ntEWS | EWS (1-355aa) | shared |
|---|---|---|
| KRT9 | RPA1 | PCMT1 |
| GNG12 | CAPRIN1 | AP2M1 |
| MPRIP | RPA2 | YTHDF2 |
| GNAI2 | PRRC2C | NCL |
| ANPEP | PRRC2A | ELAVL1 |
| KRT19 | RPA3 | RPS3 |
| LGALS1 | FAM120A | HNRNPA2B1 |
| LIMA1 | RPS6 | TAF15 |
| MYO1C | PURA | ATAD3A |
| SVIL | NUFIP2 | FUS |
| NT5E | MATR3 | HNRNPA3 |
| HSPA1B | FMR1 | RPLP2 |
| MYO1D | KHSRP | HNRNPH1 |
| | SYNCRIP | RPS5 |
| | HNRNPL | PKM |
| | HNRNPA0 | TUBA1C |
| | RTCB | HNRNPM |
| | SFPQ | TUBA1B |
| | SRSF1 | ALB |
| | | TUBB4B |
| | | PABPC1 |
| | | RPS15 |
| | | HNRNPH3 |
| | | RPL30 |
| | | VIM |
| | | TRIM21 |
| | | HIST1H4A |
| | | PGK1 |
| | | TUBB |
| | | HNRNPK |
| | | HNRNPF |
| | | ANXA2 |
| | | RPS8 |
| | | UBAP2L |
| | | HSPB1 |
| | | YWHAZ |
| | | EEF1A1 |
| | | HSPA8 |
| | | RPS18 |
| | | RPS11 |
| | | KRT7 |
| | | SMAP2 |
| | | PRDX2 |
| | | ANXA11 |
| | | RPL23 |
| | | GNAS |
| | | NPM1 |
| | | HSP90AA1 |
| | | CALM1 |
| | | KRT8 |
